# Protective immunity against malaria by a nanoparticle CIS43-based junctional vaccine alone or in combination with R21

**DOI:** 10.1101/2025.09.05.665822

**Authors:** Prabhanshu Tripathi, Ja-Hyun Koo, Xuejun Chen, Lais Da Silva Pereira, Marlon Dillon, Baoshan Zhang, Mariah Lofgren, Katelyn T. Nguyen, I-Ting Teng, Brian Bonilla, Sarah Kerscher, Wing-Pui Kong, Amy Ransier, Tyler Stephens, Yaroslav Tsybovsky, Stephanie R. Weldon, Danny C. Douek, Theodore C. Pierson, Facundo D. Batista, Azza H. Idris, Robert A. Seder, Peter D. Kwong, Tongqing Zhou

**Affiliations:** Vaccine Research Center, National Institute of Allergy and Infectious Diseases, National Institutes of Health, Bethesda, MD, USA; The Ragon Institute of Massachusetts General Hospital, Massachusetts Institute of Technology and Harvard University, Cambridge, MA, USA; Electron Microscopy Laboratory, Cancer Research Technology Program, Leidos Biomedical Research, Inc., Frederick National Laboratory for Cancer Research, Frederick, MD, USA; Department of Biology, Massachusetts Institute of Technology, Cambridge, MA 02139, USA; Department of Immunology, Harvard Medical School, Boston, MA, USA; Department of Pediatrics, Division of Pediatric Infectious Diseases & Global Health, Mass General Brigham for Children, Harvard Medical School, Boston, MA, USA; Aaron Diamond AIDS Research Center, Columbia University Vagelos College of Physicians and Surgeons and Department of Biochemistry and Molecular Biophysics, Columbia University, New York, NY 10027, USA

**Keywords:** malaria, self-assembling nanoparticle, structure-based design, vaccine, CIS43, PfCSP, R21, antibodies

## Abstract

Repetitive display of the major repeats of the *Plasmodium falciparum* circumsporozoite protein (PfCSP) is the basis for two WHO-recommended vaccines: RTS,S/AS01 and R21/Matrix-M. Recently, however, the CIS43 monoclonal antibody that preferentially targets the junctional region of PfCSP has been shown to be highly protective in humans, highlighting its junctional epitope as a key vaccine target. Here, we develop a vaccine based on the tandem repeats of the junctional epitope displayed on a self-assembling nanoparticle, and compare this CIS43-based junctional vaccine alone or in combination with the benchmark R21 vaccine, using both B cell analysis and monoclonal antibody isolation to define targeting of the immune response. Comparable reduction in liver burden was observed following vaccination with junctional and R21 vaccines at a dose of 1 μg. At a dose of 0.25 μg, a modest reduction of malaria-liver burden with the junctional vaccine was observed compared to R21. Further, combining junctional and R21 vaccines induced modestly enhanced protection compared to either vaccine alone. While the R21 vaccine elicited antibodies primarily against the major repeats, the junctional vaccine elicited antibodies against both junctional and major repeat regions. *In vivo*-B cell analysis and isolation of monoclonal antibodies confirmed differences in vaccine-induced antibody specificities. Altogether, these data suggest the nanoparticle-formatted tandem-repeated CIS43-junctional vaccine to be a promising approach to broaden immunity against malaria, either as a standalone intervention or in combination with R21.

**HIGHLIGHTS:**

- Developed a self-assembling nanoparticle-displayed junctional vaccine of PfCSP based on tandem repeats of the epitope preferentially targeted by the highly protective CIS43 antibody
- The CIS43-based junctional vaccine at low doses significantly reduced liver burden following malaria challenge in mice
- Following either low or high doses of the junctional vaccine in naïve mice, adoptively transferred B cells expressing the CIS43 inferred germline sequence yielded a high frequency of germinal center and ASC responses
- The CIS43-based junctional vaccine elicits antibodies against junctional and major repeat regions whereas the R21 vaccine elicits responses primarily against the major repeat region
- At low dose, the CIS43-based junctional vaccine given together with the R21 vaccine showed modestly improved control of liver burden compared to either vaccine alone

## INTRODUCTION

Malaria, an infectious disease caused by parasites, continues to be a major global health issue with over 263 million new malaria cases and 597 thousand deaths globally in 2023^1^. Infection is initiated when sporozoites are injected into the skin and blood by an infected mosquito. A few hundred sporozoites then quickly travel to the liver where they undergo significant expansion over the course of seven days and then are released into the blood to initiate clinical infection. Thus, prevention of sporozoites reaching the liver or infecting hepatocytes would prevent malaria infection and clinical illness. These sporozoites express a large number of surface proteins, many of which have been targeted for vaccine development. The *Plasmodium falciparum* circumsporozoite protein (PfCSP) densely covers the surface of the sporozoites and is essential for parasite development^2,3^. PfCSP has thus been the major vaccine target for vaccine development^4^.

Two WHO-approved malaria vaccines - RTS,S/AS01 ^5^ and R21/Matrix-M^6^ – display a truncated fragment of PfCSP compromised of 19 NANP repeats and the C-terminal region expressed with the hepatitis B surface antigen virus-like particle (HBsAg VLP). Four doses of RTS,S/AS01_E_ vaccine reduced cases of clinical malaria by 36% over 48 months in phase III trial^7^ and a 13% reduction in childhood mortality was reported in phase IV by WHO^8^. Three doses of 5 μg R21/Matrix-M provided 75% efficacy over a period of 12 months in phase III trial^9^. Protection by both RTS,S and R21 vaccines is largely associated with the levels of antibodies against the major NANP repeat region^9,10,11^. The deployment of these vaccines is a significant achievement and could play a major role in reducing the disease mortality. However, for both RTS,S/AS01_E_^5^ and R21/Matrix-M^6^ immunity and protective efficacy wanes over time^9,12,13^. Moreover, protection by these vaccines may be lower in perennial, high transmission areas^14^. Thus, there is still a need to improve the efficacy and durability of the current vaccines.

One approach to increase the durability of protection is to incorporate additional protective epitopes of PfCSP into a vaccine to expand the breadth of antibody responses beyond the current approved vaccines which primarily target the major repeat regions. Over the past several years, several CSP-specific human monoclonal antibodies have been isolated that identified new sites of neutralization. CIS43 preferentially targets the junctional epitope while L9 preferentially targets the NPNV minor epitope of PfCSP. The importance of these epitopes for protection is substantiated by the demonstration that a single infusion of CIS43LS was shown to be 88% efficacious over 6 months in adults in Mali^15^. Similarly, L9LS had an efficacy of 77% against clinical malaria with single subcutaneous dose of 300 mg in phase II Mali trial^16^ in 6–10 year olds. Thus, incorporating these new neutralizing epitopes not contained in RTS,S/AS01 ^5^ and R21/Matrix-M^6^ using structured-based vaccine design is a promosing approach to expanding the breadth of humoral immunity.

Self-assembling protein nanoparticle platforms (SANP) have been widely used as scaffolds for antigen display. The multivalent display of antigens provides avidity for efficient B cell receptor (BCR) crosslinking and elicitation of a strong immune response^17,18^. The inherent thermal and chemical stability provides an additional advantage for vaccine therapeutics. Lumazine synthase (LuS) from thermophilic bacterium Aquifex aeolicus self-assembles into a 60-mer icosahedral nanoparticle with a diameter of ∼150 Å. This platform was used to display the engineered outer domain of HIV-1 gp120 (eOD-GT8) and was safe when used in human clinical trial^19^. Ferritin from Helicobacter pylori forms a 24-mer octahedral nanoparticle and has been used for display of trimeric antigens like influenza hemagluttinin (HA)^20,21^, SARS coronavirus (CoV) spike^22,23^, respiratory syncytial virus (RSV) fusion glycoprotein (F)^24,25^ among others. Nanoparticle-based malaria vaccines have been reported that display the CIS43 epitope on VLP^26,27^, L9 epitope on VLP^28^, and epitopes of CIS43, L9, and MAM01 on ferritin^29^. These vaccines confer varying levels of protection in malaria mouse models. However, these vaccines have not been compared head-to-head with either RTS,S or R21 to quantitate their level of efficacy against the currently approved vaccines which will be important for further clinical development. Furthermore, developing vaccines that can mediate protection at very low doses will be critical for reducing the cost of the vaccine which is a major factor for widespread use. Last, it is important to assess whether combining vaccines to different regions of PfCSP with R21 can improve immunogenicity or protection.

Here, we utilized a self-assembling nanoparticle platform to display the CIS43 epitope, tested the protective efficacy in a mouse malaria challenge mode, and utilized adoptive B-cell transfer along with antigenic analysis of isolated antibodies to provide immunologic insight. Notably, vaccination at very low dose with tandem display of the CIS43 epitope on LuS along with the addition of a pan DR T helper epitope (PADRE) lead to protective immunity from malaria, as indicated by significantly decreased malaria-liver burden in vaccinated mice.

## RESULTS

### Design and characterization of nanoparticle immunogens incorporating the CIS43 epitope

The epitope recognized by the highly protective monoclonal antibody CIS43 encompasses a unique junctional epitope of residues NPDPNANPNVDPN as shown as a sequence schematic in **Fig. 1a** and as residue-level structure in **Fig. 1b**. As tandem repeats and self-assembling nanoparticles have been shown to be a useful approach to enhance immunogencity^30,31^, we designed multivalent immunogens based on the CIS43 epitope (**Supplementary Table S1, Supplementary Fig. 1**). Lumazine synthase 2-tandem (PADRE-LuS-2T-NPDP19) comprised a N-terminal PADRE to enhance CD4 T-cell help, the self-assembling unit of LuS and two repeats of the CIS43 epitope with KQPADG added at the N-terminus and with a ggsgg spacer (**Fig. 1c, Supplementary Table S1**). Lumazine synthase 3-tandem (PADRE-LuS-3T-NPDP19) comprised the same as PADRE-LuS-2T-NPDP19 with an additional tandem repeat (**Fig. 1c, Supplementary Table S1**). For simplicity of nomenclature in the rest of the manuscript, the NPDP19 is omitted from double and triple tandem constructs and these are referred as PADRE-LuS-2T or 3T. To determine if the orientation of the PADRE helper epitope influenced immunity and protection, constructs containing PADRE at the C-terminal with the junctional epitope at the N-terminal of LuS were designed and are referred as 2T-LuS-PADRE or 3T-LuS-PADRE for 2 and 3-tandem repeats, respectively. Expression and purification of the constructs yielded a single major peak with protein eluting at ∼12 minutes (corresponding to molecular weight of 26 and 32 kDa); SDS-PAGE analysis revealed each of the major peaks to comprise a single band (**Fig. 1d, Supplementary Fig. 1**). Negative stain electron microscopy (EM) showed assembled nanoparticles with ∼160 Å diameter (**Fig. 1e**), and the antibody binding revealed strong recognition for PfCSP-directed junctional monoclonal antibodies including CIS43^32^, D3^33^, and P3-43^34^ as well as L9^35^ (targeting the minor repeat region), and 317^36^ (targeting the major repeat region). Strong recognition was also observed for CIS42^32^, a non-neutralizing antibody that binds the junctional region (**Fig. 1f**). Thus, while developing vaccines displaying only the junctional epitopes of PfCSP, there is broad recognition by monoclonal antibodies that also target the major and minor repeats.

**Figure 1.**
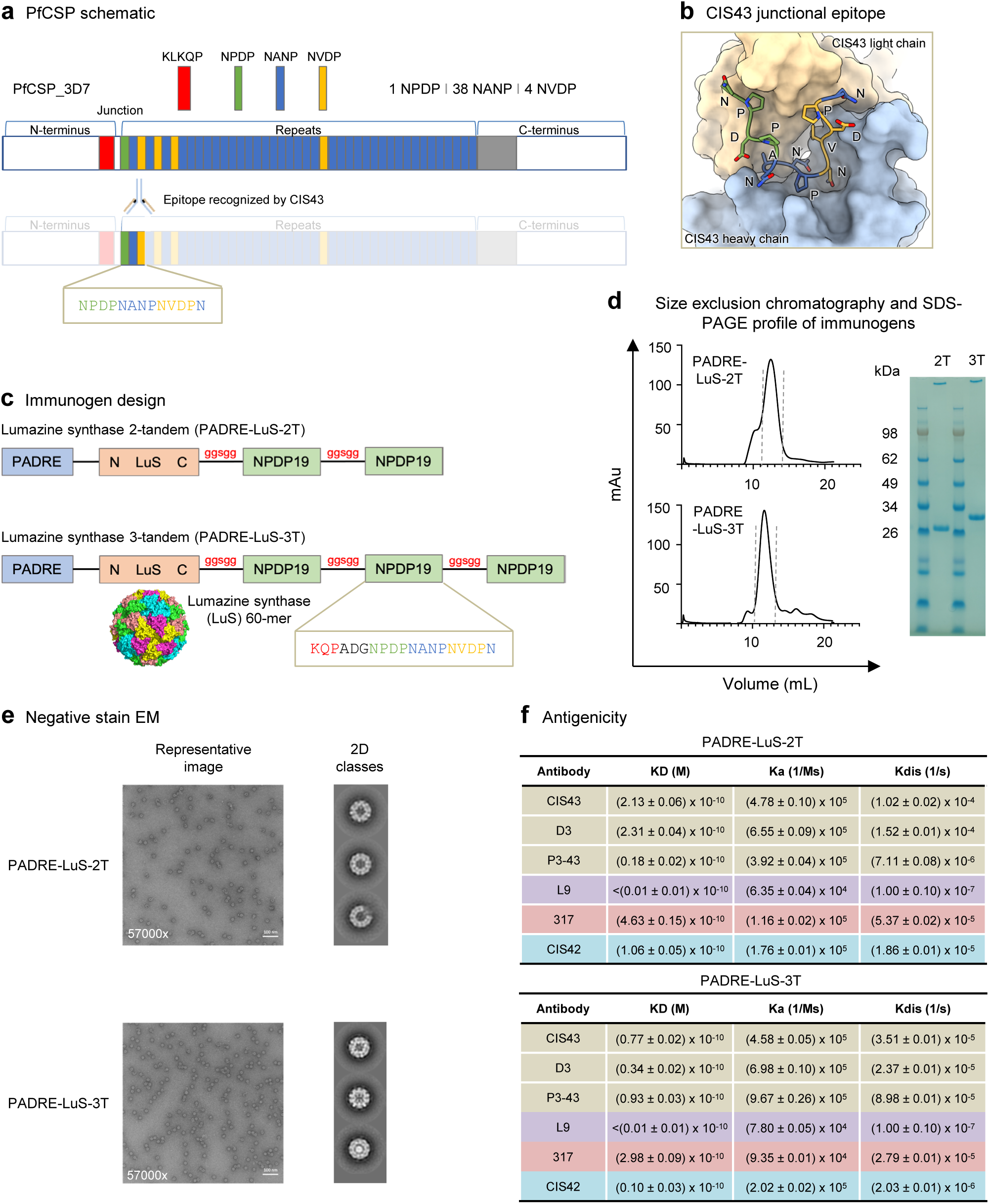
Design and characterization of nanoparticle immunogens incorporating the CIS43 epitope. **a** Schematic of plasmodium falciparum circumsporozoite protein (PfCSP) highlighting the CIS43 epitope. **b** Structure of CIS43 bound to peptide 21 of PfCSP. **c** Design of nanoparticle immunogens. **d** Characterization of immunogens by size exclusion chromatography and SDS-PAGE. **e** Negative stain EM shows properly assembled nanoparticle immunogens. **f** Antigenicity of nanoparticle immunogens to junction, minor, and major-directed antibodies assessed by biolayer interferometry.

### Tandem-displayed CIS43 epitopes on LuS nanoparticle induce protective anti-malarial responses

To test if the tandem display of junctional epitope increases the efficacy of the designed immunogens, mice were immunized with a single (NPDP19-LuS-PADRE) and double tandem repeat (2T-LuS-PADRE) of the CIS43 epitope displayed on lumazine synthase. These nanoparticle vaccines were compared to a single NPDP19 epitope linked to KLH. These junctional immunogens were also compared to the R21 vaccine which primarily targets the repeat regions of PfCSP and serves as the benchmark. Mice received two immunizations four weeks apart. Sera was collected at weeks 2 and 6 and analyzed with biolayer interferometry (BLI) to assess antibody responses against PfCSP, major epitope (NANP9), and junctional epitope (NPDP19). At week 2, R21-immunized group elicited high responses against PfCSP, NANP_9_, and almost negligible responses against NPDP19, as expected since the vaccine does not contain the junctional epitope. In contrast, only the 2T-LuS-PADRE group showed measurable responses against NPDP19 at week 2. At week 6, the R21 elicited responses were at similar levels as week 2; however, all the other tested immunogens showed strongly enhanced elicited antibody levels. The NPDP19-KLH control and the 2T-LuS-PADRE groups had the highest responses when compared across all three probes. It was notable that an inverse dose response trend was observed for the NPDP19-LuS-PADRE groups with the lower dose of 1 μg elicited higher antibody responses compared to the 10 μg dose, however these differences were not significant. Morever, the 2T-LuS-PADRE group showed improved antibody responses versus the NPDP19-LuS-PADRE group (**Fig. 2b**).

**Figure 2.**
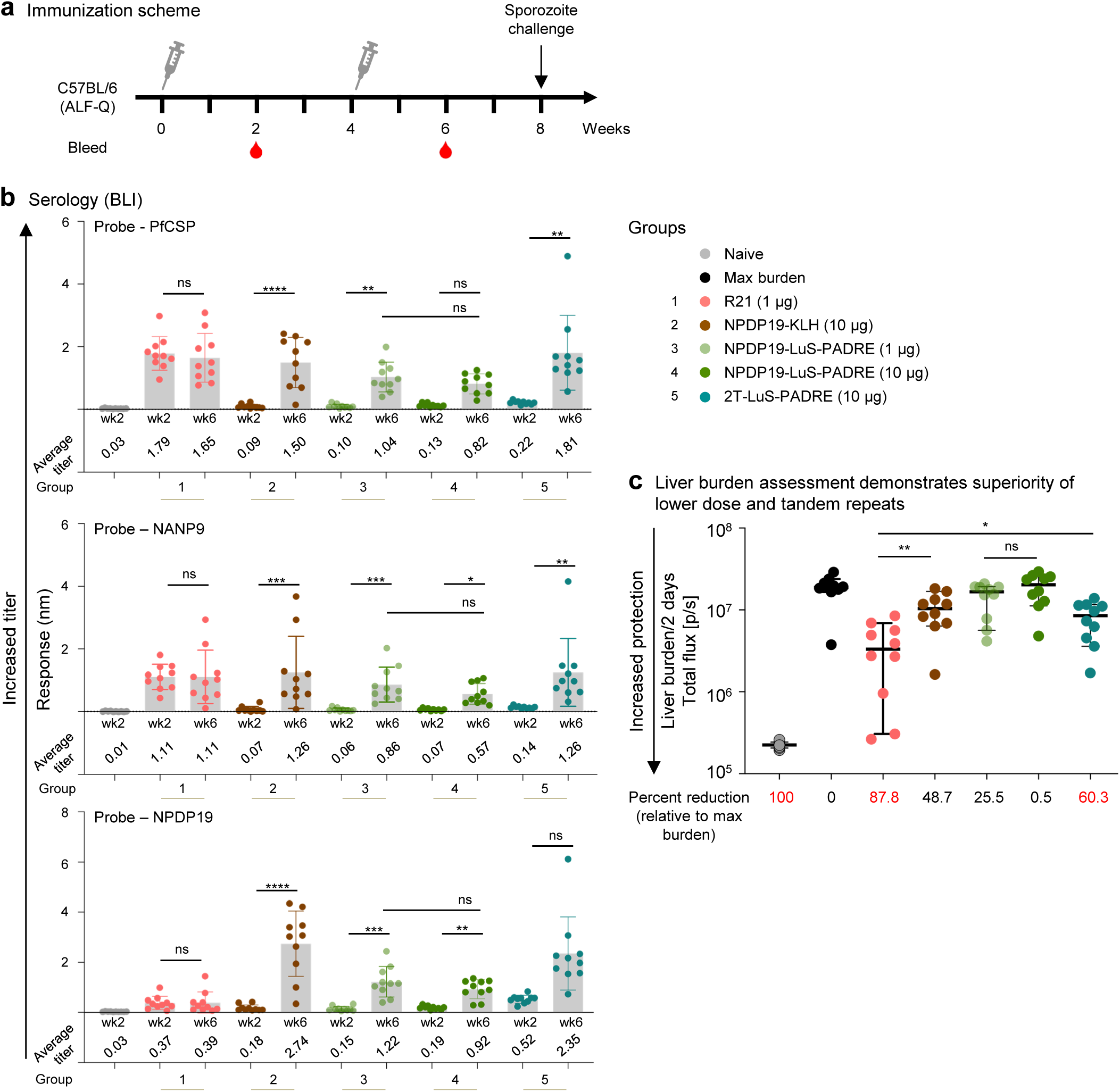
Malaria protection by vaccination with tandem-displayed CIS43 epitope on LuS nanoparticle. **a** Immunization schema comprising of 2 immunizations 4 weeks apart followed by sporozoite challenge at week 8. **b** Serum antibody titers at weeks 2 and 6 assessed by biolayer interferometry (BLI) against biotinylated PfCSP, NANP9 (major repeat), and NPDP19 (junction). The average titer of each group is listed. **c** Liver burden assessment of immunized mice at day 2 post Pb-PfCSP-Luc sporozoite challenge is shown with the horizontal lines denoting the median flux; error bars show 95 % confidence interval; N=10 mice per group. The percent reduction is calculated based on geometric mean of each group and values above 50 % are highlighted in red. * p < 0.05, ** p < 0.01 as calculated by two-tailed Mann–Whitney test.

Vaccine efficacy was assessed by calculating the reduction in liver burden at day 2 after challenge with a well-established model using chimeric Pb-PfCSP-Luc sporozoites. The mice that received the R21 vaccine at 1 μg showed 87.8% reduction in liver burden (**Fig. 2c)**, whereas the 2T-LuS-PADRE group at 10 μg showed 60.3% reduction. By contrast the NPDP19-LuS-PADRE vaccine showed no significant reduction in liver burden at either the 10 or 1 μg dose compared to the maximum liver burden in infected naïve animals. Of note, the inverse dose response trend that was observed with antibody titers (**Fig. 2b**) was recapitulated in the liver burden assay, with the lower dose group (1 μg) showing ∼ 25% protection while the (10 μg) dose group had no reduction in liver burden (**Fig. 2c**). However, these differences did not reach statistical significance.

### Immunogenicity and Protection by LuS-displayed vaccine

As the 2T-LuS-PADRE showed enhanced antibody and control of liver burden compared to the single NPDP19-LuS-PADRE, we next performed a more extensive analysis using the 2T-LuS-PADRE at three different doses of 0.2, 1, and 5 μg and compared to R21 at 1 μg following the same immunization schema as the previous experiment (**Fig. 3a**). We chose this dosing range, because the current R21 is licensed for administration at 10 μg in adults^9^, with dosing calculation indicating a 1 μg dose in mouse to be equivalent to ∼244 μg in adult human, and 0.2 μg dose to be equivalent to ∼50 μg^37^.

**Figure 3.**
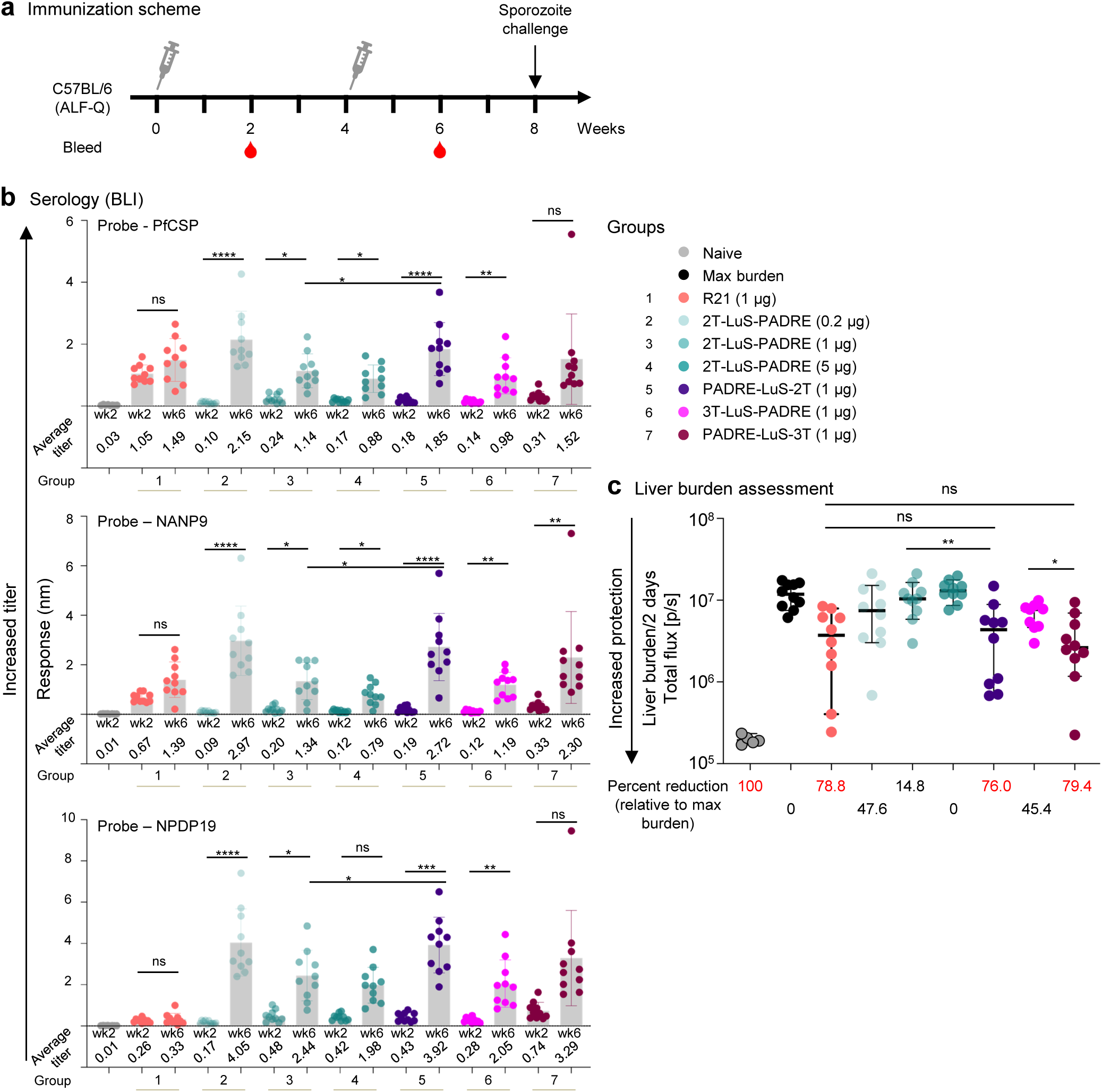
Tandem LuS-displayed CIS43 epitope vaccine provides protection equivalent to R21 at 1 ug dose. **a** Immunization schema comprising of 2 immunizations 4 weeks apart followed by sporozoite challenge at week 8. **b** Serum antibody titers at weeks 2 and 6 assessed by biolayer interferometry (BLI) against biotinylated PfCSP, NANP9 (major repeat), and NPDP19 (junction). The average titer of each group is listed. **c** Liver burden assessment of immunized mice at day 2 post Pb-PfCSP-Luc sporozoite challenge is shown with the horizontal lines denoting the median flux; error bars show 95 % confidence interval; N=10 mice per group. The percent reduction is calculated based on geometric mean of each group and values above 50 % are highlighted in red. * p < 0.05, ** p < 0.01 as calculated by two-tailed Mann–Whitney test.

All three 2T-LuS-PADRE dose groups showed significantly enhanced antibody responses at week 6 compared to week 2. Of note, there was an inverse relationship between the dose administered and antibody production with the lowest dose of 0.2 μg having the highest antibody response against all three probes: PfCSP, NANP9, NPDP19 (**Fig. 3b**). Consistent with the prior data (**Fig. 2b**), R21 showed antibody responses to only PfCSP and NANP9, and responses did not increase significantly after the second immunization (**Fig. 3b**).

To determine if additional valency of junctional epitopes would alter immunity and protection, immunogens containing 3 tandem repeats were generated. In addition, we also assessed whether placement of the added T-cell epitope (PADRE) to the N-terminus instead of the C-terminus influenced immunity and protection. At the 1 μg dose, the immunogen containing 2 tandem repeats with N-terminally placed PADRE (PADRE-LuS-2T) yielded significantly higher antibody responses than C-terminal positioned PADRE across all three probes (**Fig. 3b**, comparing groups 3 and 5 at week 6). For the 3 tandem repeat immunogen with N-terminal PADRE (PADRE-LuS-3T) the responses were significantly higher against the major repeat probe (NANP9) and showed a trend toward higher responses against PfCSP and NPDP19, but these differences were not significant (**Fig. 3b**, comparing groups 6 and 7 at week 6). PADRE-LuS-2T and PADRE-LuS-3T elicited similar antibody responses (**Fig. 3b**, comparing groups 5 and 7 at week 6).

We then assessed liver burden after sporozoite challenge at week 8. Reduction in liver burden was higher with N-terminal placed PADRE with PADRE-LuS-2T or 3T showing significantly greater reduction in liver burden at 76.0% and 79.4% compared to 14.0% and 45.4% with PADRE at the C-terminal (**Fig. 3c**). PADRE-LuS-2T or 3T had comparable reduction in liver burden than R21 (78.8%) at the same dose of 1 μg. Of note, there was an inverse dose response effect on liver burden with 2T-LuS-PADRE with ∼ 48% reduction at the 0.2 μg dose and no reduction at the 5 μg dose. These data are consistent with the inverse dose effect of this immunogen on antibody levels. In general, the protection correlated with the antibody binding, with the N-terminal PADRE-based immunogens containing either 2 or 3 NPDP19 repeats showing comparable protection to R21 at a dose of 1 μg (**Fig. 3c**).

### Tandem LuS-vaccine at 0.25 **μ**g dose provides higher protection relative to R21

Based on the results showing that the N-terminal position of PADRE and lower doses may be optimal for immunity and protection (**Figs. 2** and **3**), we assessed lower doses of PADRE-LuS immunogen containing 2 or 3 tandem repeats of NPDP19 and compared to R21 (**Fig. 4**). As observed in the prior figure with the 2T-LuS PADRE, the lower dose of PADRE-LuS-2T or 3T at 0.25 μg yielded increased antibody titers than the 1 μg dose for PfCSP, NANP9 and NPDP19 (**Fig. 4b**). By contrast, the antibody titers against NANP9 or PfCSP were lower at 0.25 μg than at 1 μg dose. (**Fig. 4b**). These data suggest that immunity and protection may in fact be optimal at lower doses of the nanoparticle junctional vaccines in contrast to the benchmark R21 vaccine which shows a standard dose response.

**Figure 4.**
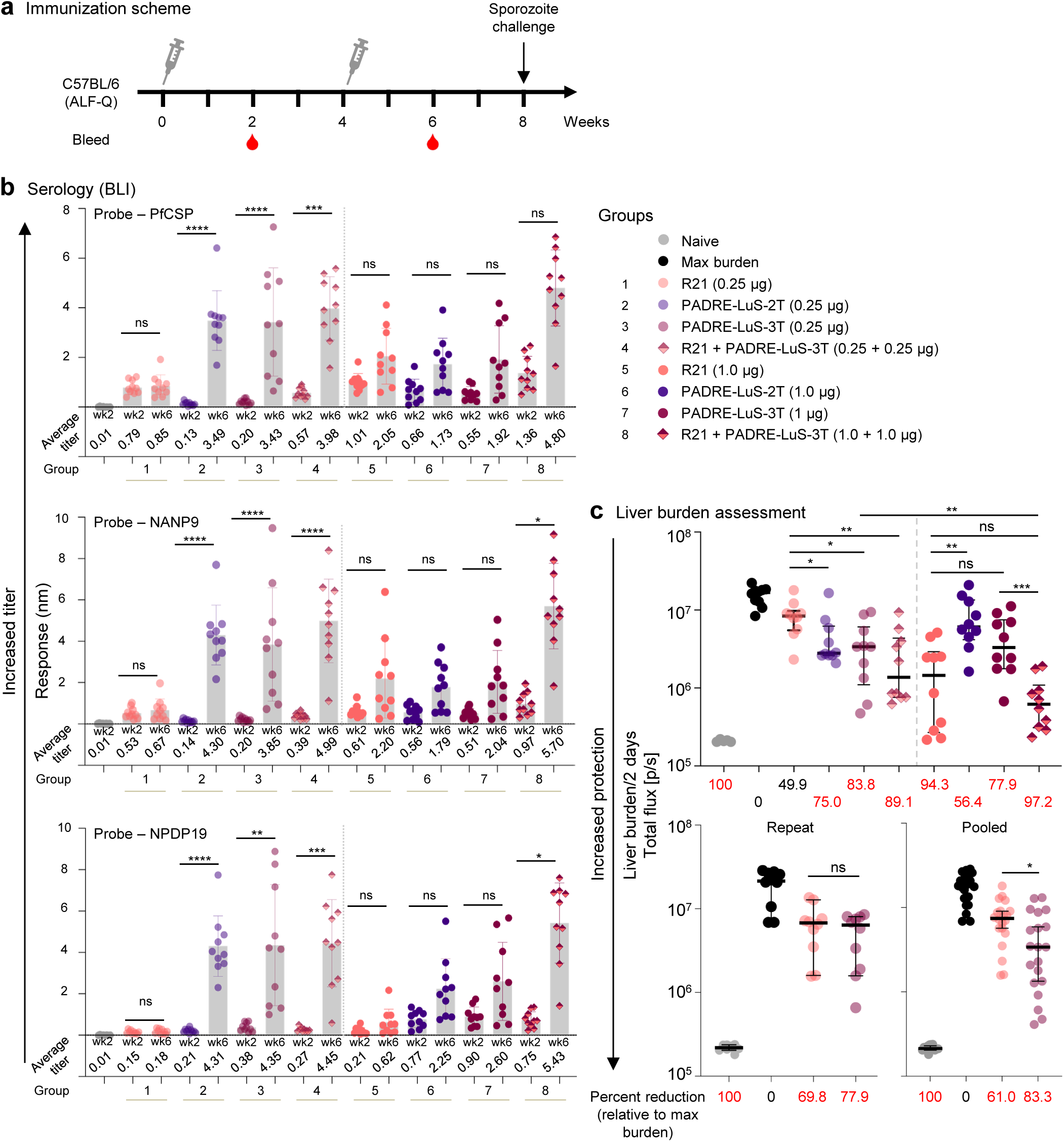
Tandem LuS-displayed CIS43 epitope vaccine provides increased protection relative to R21 at 0.25 ug dose. **a** Immunization schema comprising of 2 immunizations 4 weeks apart followed by sporozoite challenge at week 8. **b** Serum antibody titers at weeks 2 and 6 assessed by biolayer interferometry (BLI) against biotinylated PfCSP, NANP9 (major repeat), and NPDP19 (junction). The average titer of each group is listed. **c** Liver burden assessment of immunized mice at day 2 post Pb-PfCSP-Luc sporozoite challenge is shown (top panel) with the horizontal lines denoting the median flux; error bars show 95 % confidence interval; N=10 mice per group. The repeat experiment is shown for R21 and PADRE-LuS-3T at 0.25 ug; N=10 mice per group (bottom left panel). The pooled results of two independent experiments is shown with N=20 mice per group (bottom right panel). The percent reduction is calculated based on geometric mean of each group and values above 50 % are highlighted in red. * p < 0.05, ** p < 0.01 as calculated by two-tailed Mann–Whitney test.

To next assess protection following the different doses of the PADRE-LuS-2T or 3T vaccines, in the first experiment (**Fig. 4c** upper panel) both the PADRE-LuS-2T and 3T immunogens at 0.25 μg dose showed 75.0% and 83.8% reduction of liver burden respectively, which were significantly better than the protection of 49.9% from R21 at 0.25 μg dose. However, at 1 μg dose, R21 had 94.3% reduction of liver burden which was significantly better than the PADRE-LuS-2T at 56.4%. The PADRE-LuS-3T at a dose of 1 μg had 77.9% reduction of liver burden which was not significantly different than R21 (**Fig. 4c** top right panel). In a second experiment, we compared R21 and PADRE-LuS-3T at the 0.25 μg dose (**Fig. 4c**, bottom left).

Both vaccines showed comparable reduction in liver burden. Data from two independent experiments were pooled and with N=20 mice/group, PADRE-LuS-3T had significantly higher protection of 83.3% compared to R21 with 61% protection at 0.25 μg dose (**Fig. 4c**, bottom right). Analysis of antibody titers and protection showed significant correlation for all three probes with the highest correlation for PfCSP (R^2^ = 0.3776) and NANP9 (R^2^ = 0.3671), and lower correlation with NPDP19 (R^2^ = 0.2119) (**Supplementary Fig. 2**).

Although the junctional vaccines were able to elicit broad PfCSP, NANP9, and NPDP19 responses, we assessed whether combining them with R21, which primarily elicits NANP9 resposes would improve immunity and protection. Indeed, the highest and broadest antibody responses were observed by combining the vaccines (**Fig. 4b**). Moreover, the reduction in liver burden was 89.1% in mice that received both vaccines at 0.25 μg dose compared to 49.9% (R21), and 83.8% (PADRE-LuS-3T) alone. At the 1 μg dose, reduction in liver burden was 97.2% in mice that received both vaccines and 77.9% (PADRE-LuS-3T) and 94.3% (R21) alone. Thus, there is a modest but significant additive effect of having the junctional and R21 vaccines mixed (**Fig. 4c** bottom panel) at the lowest doses of both vaccines.

### Frequency of germinal center B cells and antibody secreting cells following junctional vaccination

Based on the enhanced antibody responses and protection by the lower doses of the junctional vaccines, the analysis was extended to assess the in vivo mechanism of priming through the the germinal center (GC) response. Inferred germline CIS43 (iGL_CIS43) B cells that respond to the NPDP19 junctional epitope were used to quantitate the response in vivo to vaccination. We adoptively transferred CD45.2^+^ NPDP19-specific inferred germline CIS43 (iGL_CIS43) B cells into CD45.1+ host mice and then immunized them with either a low (0.25 μg) or high (10 μg) dose of PADRE-LuS-3T (**Fig. 5a**). Mice were immunized with NPDP19-KLH (positive control) as it was shown to induce high GC response and effective elicitation of CIS43 germline-precursor B cells in a previous study^33^ or PADRE-LuS-3T-2NPNV (a minor repeat immunogen) as a negative control for specifity. GC responses were assessed 10 days post-immunization.

**Figure 5.**
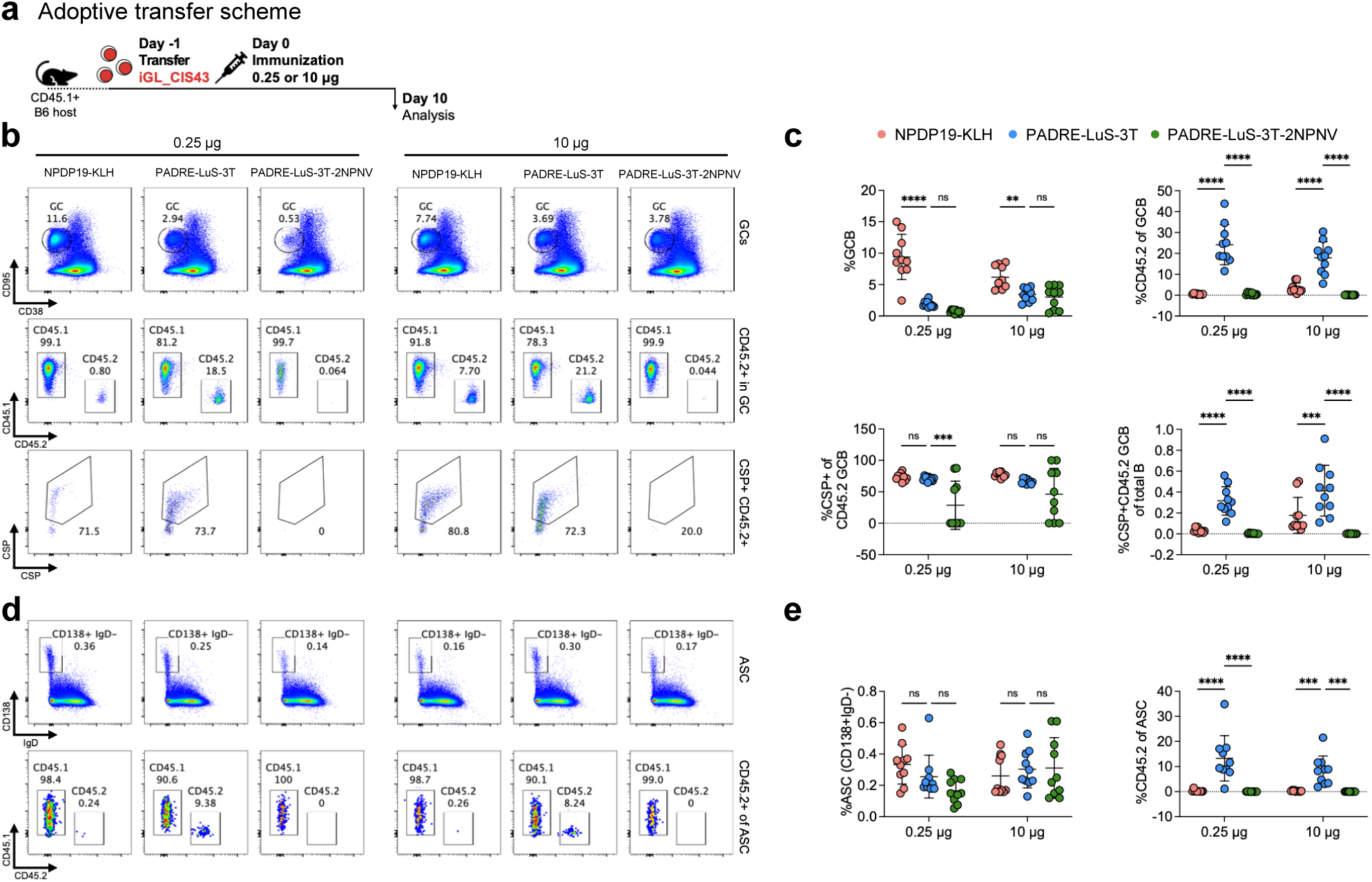
PADRE-LuS-3T elicits robust germinal center response in CIS43 KI-mice. **a** Schematic of adoptive transfer model to evaluate iGL_CIS43 responses to PADRE-LuS-3T immunogen. **b** Representative FACS plot of GC response and CD45.2+ B cells in GC. **c** Quantification of GC formation, CD45.2+ in GC, CSP+ in CD45.2+GCB and CSP+CD45.2+GCB frequency in total B cells. N = 10 mice in each group. The data is pooled from two independent experiments. Each symbol represents a different mouse. One-way ANOVA was applied, and the bars indicate mean ± SD. p>0.05 represents no significance (ns), *p<0.05, **p<0.01, ***p<0.001, ****p<0.0001. d Representative FACS plot of antibody secreting cell (ASC) response and CD45.2+ B cells in ASC. e Quantification of ASC frequency, and CD45.2+ of ASC. N = 10 mice in each group. The data is pooled from two independent experiments. Each symbol represents a different mouse. One-way ANOVA was applied, and the bars indicate mean ± SD. p>0.05 represents no significance (ns), *p<0.05, **p<0.01, ***p<0.001, ****p<0.0001.

In the NPDP19-KLH group, the CD45.2+ B cell ratio in the GC was 7.70 in the high-dose group (10 μg) compared to 0.80 in low-dose group (0.25 μg), resulting in a higher frequency of CD45.2+ GC B cells among total B cells (**Figs. 5b** and **5c**). However, in the PADRE-LuS-3T group, the low dose induced a GC response of iGL_CIS43 B cells comparable to the high-dose group. Furthermore, the GC response from both the low and high doses of PADRE-LuS-3T was significantly higher than the response from NPDP19-KLH, suggesting that the tandem immunogen could induce an immune response more efficiently than KLH conjugates. As expected, PADRE-LuS-3T-2NPNV did not induce any responses.

The ability of the PADRE-LuS-3T vaccine to efficiently expand CIS43 lineage B cells which induced broad antibody responses and protection shows the effectiveness of this structure guided nanoparticle approach to the junctional epitope. Thus, the next focus was to determine whether CIS43 type antibodies were generated. A restricted set of “Core8” mutations; four on the heavy chain (M34I_H_, N52K_H_, K58R_H_ and V98I_H_) and four on the light chain (S27_A_N_L_, V27_B_I_L_, Q89H_L_ and T94S_L_) have been identified to correlate highly with malaria protection for CIS43 variants^33^. To assess if immunization by PADRE-LuS-3T elicited these key CIS43-like mutations, paired sequence analysis was performed at day 24 post immunization (**Supplementary Fig. 3a**). Hotspot analysis revealed elicitation of critical “Core8” mutations with both 0.25 μg and 10 μg of PADRE-LuS-3T (**Supplementary Fig. 3b-d**). Thus, CIS43-like sequence features can be recapitulated by immunization with PADRE-LuS-3T.

We also examined the differentiation of antibody-secreting cells (ASCs) of iGL_CIS43. No significant differences in the frequency of class-switched ASCs (CD138+IgD-) among total B cells were observed between NPDP19-KLH, PADRE-LuS-3T, and PADRE-LuS-3T-2NPNV immunization groups (**Figs. 5d** and **5e**). However, significant ASC differentiation of iGL_CIS43 was specifically observed in the PADRE-LuS-3T group. Furthermore, there was no significant difference in the ASC response between the high- and low-dose groups. Thus, the PADRE-LuS-3T immunogen elicited robust GC and ASC responses, with the low dose inducing responses as strong as those observed with the high dose.

### Antibody profiling following junctional or R21 vaccination

To isolate the vaccine elicited protective anti-malaria antibodies and compare the antibodies elicited by PADRE-LuS-3T and R21 vaccines, we selected mice with the lowest liver burden post challenge for B cell sorting and antibody identification. Two mice from the PADRE-LuS-3T (0.25 μg)-immunized group (mice 5147 and 5149) and one from the R21 (1 μg) - immunized group (mouse 5116) were analyzed. Index-sorted IgG+ B cells that are triple positive for the full-length PfCSP, the major repeat peptide (NANPNANPNANPNAN; P29) and the junctional peptide (NPDPNANPNVDPNAN; P21) probes were analyzed from spleens of vaccinated animals (**Supplementary Fig. 4**). Based on the flow cytometry analysis, the R21-immunized mouse elicited mainly P29-specific PfCSP+ memory B cells and very few P21-reactive PfCSP+ memory B cells; in contrast, the PADRE-LuS-3T immunized mice showed much higher percentages of PfCSP+ memory B cells that also bound P21 and/or P29 (**Fig. 6a**). These results suggest that PADRE-LuS-3T elicited more cross-reactive IgG+ B cells that can target both the junctional and major repeat peptides of PfCSP whereas R21 elicited predominantly the major repeat peptide specific IgG+ B cells with few targeting the junctional peptide. Consistent with the B cell antigenicity analysis results, the ELISA screening of the supernatants of antigen sorted B cells against PfCSP, junctional peptide P21 and major repeat peptide P29 confirmed that PADRE-LuS-3T immunized mice 5147 and 5149 expressed more cross-reactive (PfCSP+/P21+, PfCSP+/P29+ or PfCSP+/P21+/P29+) antibodies than the R21-immunized mouse 5116 (**Fig. 6b**). There was only one strong PfCSP-binding antibody, one strong P29-binding antibody and zero P21+/PfCSP+ antibody identified from 72 antigen-sorted B cells of R21-immunized mouse 5116, consistent with the extremely low frequency of P21+/PfCSP+ B cells in this mouse (**Figs. 6b, c**). In comparison, 29 of the 78 mouse 5149 supernatants and 2 of the 84 mouse 5147 supernatants are triple positive in binding to PfCSP, P21 and P29. Moreover, mouse 5147 also have 6 strong PfCSP binders and a P21/P29 double binder. Antibody sequence analysis revealed a large clonal family (green highlighted in **Fig. 6c**) of PfCSP/P21/P29 triple positive antibodies originated from two different PADRE-LuS-3T immunized mice. They all have a IGHV1-72*01 heavy chain with a 10 aa CDRH3 and a IGKV5-43*01 or IGKV5-48*01 light chain with a 9 aa CDRL3. There is also another clonal family of antibodies found in mouse 5147 that bind strongly only to PfCSP but not to the junctional or major repeat peptide (blue highlighted in **Fig. 6c**). The B cell antigenic analysis and the isolated antibodies demonstrate that the PADRE-LuS-3T elicited more triple positive antibodies that bind both junctional and major repeat peptides than R21, which failed to elicit junctional peptide specific antibodies.

**Figure 6.**
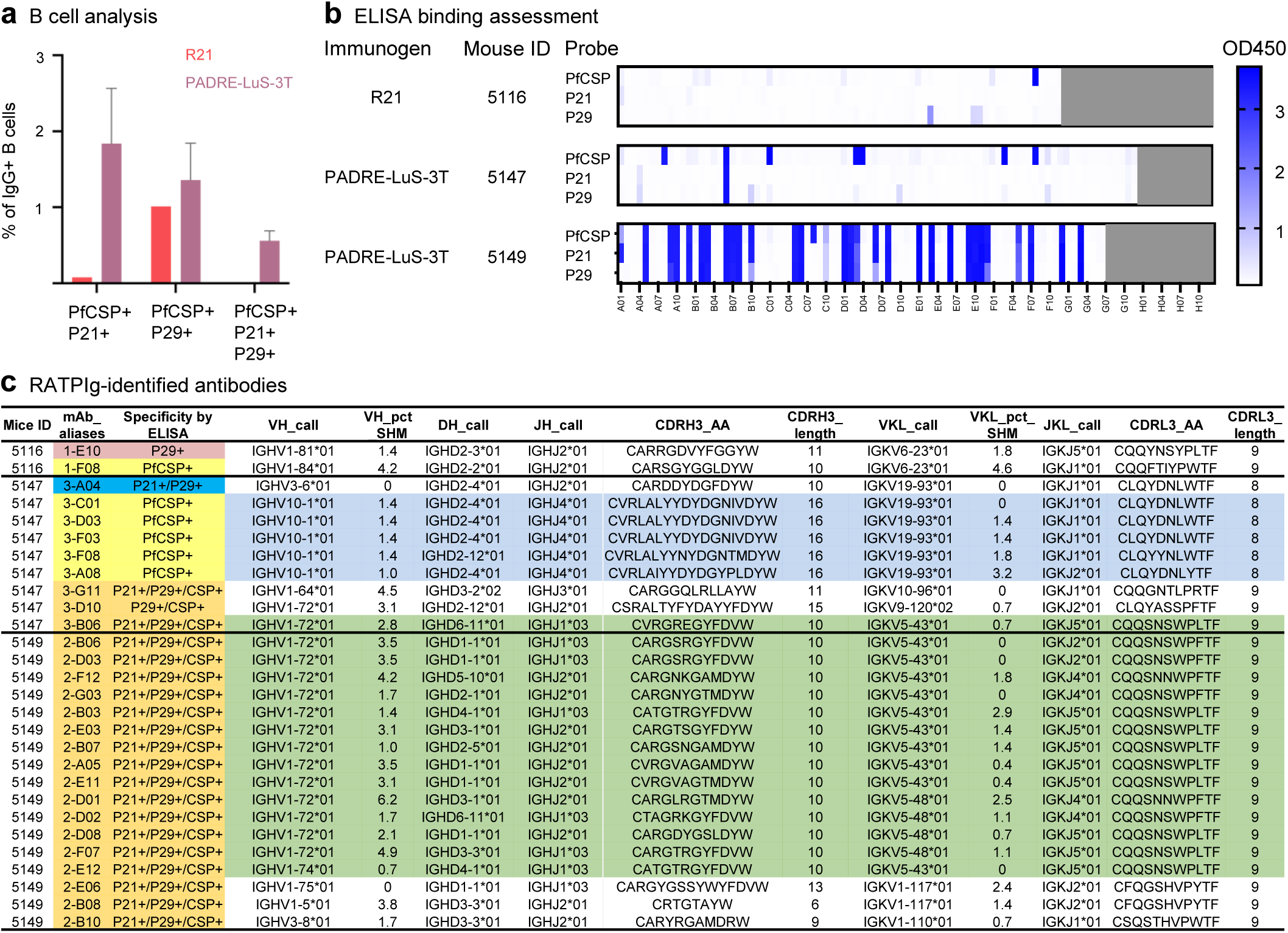
B-cell sorting and antibody antigenic analysis reveal PADRE-LuS-3T elicits cross-reactive responses in contrast to R21 in mice. **a** Percentages of PfCSP+/P21+, PfCSP+/P29+ and PfCSP+/P21+/P29+ (+++) B cells among all IgG+ B cells in an R21-immunized mouse 5116 (black dot) and two PADRE-LuS-3T immunized mice 5147 and 5149 (red triangle), revealed by flowcytometry. **b** ELISA screening of RATPIg supernatants in 96-well plates containing expressed IgGs from R21 or PADRE-LuS-3T immunized animals. Each well corresponds to an index sorted B cell except the gray-shaded wells (no cell). The binding (OD450) of the supernatants to PfCSP, P21 and P29 was plotted in heatmaps. **c** List of antigen-specific antibodies identified by RATPIg from sorted splenic B cells of R21 (mouse 5116)- or PADRE-LuS-3T (mice 5147 and 5149)-immunized animals. Antibodies of similar clonal families were highlighted in blue or green color.

## DISCUSSION

In this study, we designed immunogens that displayed the junctional region of PfCSP on self-assembling LuS nanoparticle platform. The genetic fusion of the junctional peptide NPDP19 to LuS eliminated the need of covalent coupling and enabled the production of immunogens in a single component fashion. The added glycans on LuS surface facilitate a tag-free purification using lectin affinity chromatography. The immunogens were well assembled as shown by electron microscopy and bound tightly to known malaria antibodies. The immunogens were evaluated for their malaria protective efficacy in an in vivo mouse model of malaria and benchmarked to the best-in-class malaria vaccine, R21. The PADRE-LuS-3T immunogen was comparable in protection to the R21 vaccine with some data supporting modestly enhanced protection at a very low dose (0.25 μg). When combined with R21, an additive trend in protective efficacy was observed which further highlighted the importance of junctional epitope in the design of PfCSP-based malaria vaccines.

Critical features of the most effective immunogen designed in this study are the tandem display of the NPDP19 peptide and where the placement of PADRE helper epitope occurred. The junctional peptide when displayed in tandem (2T-LuS-PADRE) enhanced protective efficacy to 60.3% compared to no protection with a single repeat (NPDP19-LuS-PADRE) at 10 μg dose. The C terminus of CSP (C-CSP) is known to contain T helper epitopes^38,39,40^, which is part of the RTS,S and R21 vaccines. In the absence of the C-CSP, a PADRE peptide was included to provide additional T cell help. PADRE has been used in vaccines for activation of CD4+ T cells to boost immune responses^41^ and was shown to be safe and clinically feasible in humans in a cytomegalovirus (CMV) peptide vaccine study^42^. Interestingly, addition of PADRE to the N-term of LuS provided significantly better protection than the C-term PADRE. The mechanism by which the N-terminal location of PADRE impacted immunogenicity was not assessed in these studies.

A key feature of this study was to utilize the well established model of adoptive transfer of B cells expressing the specific antibody sequence being targeted to assess how the vaccines activate B cells and if this correlated with antibody responses. Mice with B cells expressing the pre-rearranged inferred germline sequence of protective human antibodies, in this instance CIS43, provide a platform for the recapitulation of the in vivo affinity maturation process by which those protective human antibodies first arose, and variations to that ontogeny arising during the replay may be in themselves of biological or therapeutic interest^33^. The adoptive transfer aspect further enhances model verisimilitude, as the frequency of specific B cell precursors within the population can be an important determinant of immunogenicity^43^. B-cell antigenic analyses revealed broad responses elicited by PADRE-LuS-3T, whereas, R21-elicited responses were mainly directed to the major repeat region of CSP. Importantly, a recent analysis of the breadth of PfCSP responses following R21 in malaria-naïve adults showed a heavily skewed repertoire dominated by NANP-specific IGHV3-30/3-33 lineages^44^. Despite the lack of junctional region in R21, these antibodies display cross-reactivity to the junctional region of CSP because of epitope similarity. CSP-targeting antibodies that cross-react to junction (NPDP), minor (NVDP), and major (NANP) regions are often associated with high affinity and potency^45,46,47^. The isolation of such cross-reactive antibodies from mice immunized with PADRE-LuS-3T underscores it’s potential advantages as elicitation of broad responses may translate into improved protective efficacy.

A notable finding was the inverse dose effect observed with the LuS-based immunogens. Antibody responses and protective efficacy were higher at very low doses. This inverse dose response is in agreement with a prior study, which showed that the highest doses of RTS,S and R21 showed lower protection^48^. This apparent paradox may be explained by abundant antigen relaxing the competition for T cell help; this intra-GC competition is key to driving affinity maturation^49^. In the case of R21, the number of redundant major repeat epitopes presented may further functionally relax this pressure, saturating the GC without meaningfully selecting for high affinity clones. The in vivo immunization model demonstrates that in contrast, even at low doses, LuS-based immunogens presenting the non-redundant PfCSP junctional region can drive productive engagement. Intact, rather than degraded, antigen retention may also influence these GC dynamics: a recent study using an LuS-based HIV immunogen, eOD-60mer, found that a high proportion of the antigen remained intact in the follicles out to seven days after immunization^50^, though whether the immunogen delivered here displays similar structural stability at this critical anatomical site is unknown. Sufficient, but not overwhelming, site-specific availability of intact antigen during the initial GC response may thus modulate not only the magnitude but also the protective quality of the antibody response.

It is difficult to compare human and mouse dosing, but based on human equivalent dose calculation, the 0.25 μg dose/mouse is 6–12-fold higher than the amount administered to children-adults, so there is the potential for having a very low dose vaccine for humans which would be important to reduce cost. Further studies in NHP with a wide dose range to compare the immunogenicity and durability of the junctional vaccine and R21 will further inform clinical development. The data shown here substantiates the junctional epitope as a key site for sporozoite-stage anti-malaria antibodies and protection. Moreover, the demonstration that combining the junctional and R21 vaccines may be optimal for magnitude, breadth of antibody responses, and better protection suggests this as a potentially improved approach in humans. Future studies combining junctional vaccines with R21 together or in sequence will be evaluated for improvement over either approach alone to establish breath and efficacy. Further incorporation of blood-stage vaccines with the junctional vaccine could enable development of a multi-stage malaria vaccine.

## METHODS

### Protein production and purification

For protein expression, 3 ml of Turbo293 transfection reagent (Speed BioSystems) was mixed with 50 ml Opti-MEM medium (Life Technology) and incubated at room temperature (RT) for 5 min. 1 mg plasmid DNAs was mixed with 50 ml of Opti-MEM medium in a separate tube, and the mixture added to the Turbo293 Opti-MEM mixture. The transfection mixture was incubated for 15 min at RT then added to 800 ml of Expi293 cells (Life Technology) at 2.5 million cells/ml. The transfected cells were incubated overnight in a shaker incubator at 9% CO2, 37 °C, and 120 rpm. On the second day, about 100 ml of Expi293 expression medium was added. On day 6 post transfection, supernatants were harvested, filtered. Proteins were purified from the supernatant using GNA lectin affinity chromatography. GNA resin was equilibrated with 10 column volumes (CVs) of PBS (pH 7.4) followed by supernatant loading. Unbound protein was removed by washing with 20 CVs of PBS buffer. Bound protein was eluted with 250 mM methyl α-d-mannopyranoside in PBS buffer (pH 7.4). All proteins were purified by size exclusion chromatography on Superdex 200 Increase 10/300 GL in PBS and further characterized by SDS-PAGE.

### Negative-stain electron microscopy (EM)

Samples were diluted to 0.02–0.05 mg/ml with a buffer containing 10 mM HEPES, pH 7, and 150 mM NaCl. A 4.7-µl drop of the diluted sample was applied to a glow-discharged carbon-coated copper grid for approximately 15 s. The drop was removed using blotting paper, and the grid was washed three times with 4.7-µl drops of the same buffer. Adsorbed proteins were negatively stained by applying consecutively three 4.7-µl drops of 0.75% uranyl formate and removing each drop with filer paper. Micrographs were collected using SerialEM45 on an FEI Tecnai T20 electron microscope operated at 200 kV and equipped with an Eagle CCD camera or using EPU on a ThermoFisher Talos F200C electron microscope operated at 200 kV and equipped with a Ceta CCD camera.

### Antigenicity characterization by Biolayer Interferometry

The antigenicity of nanoparticles was assessed by binding to junctional, minor, and major directed antibodies. Antibodies were diluted to 5 ug/mL in (PBS + 1% BSA), immobilized on AHC biosensors to a density of 0.1 nm, and dipped into varying nanoparticle concentrations. Assays were performed with agitation at 30°C. In all Octet measurements, parallel correction to subtract systematic baseline drift was carried out by subtracting the measurements recorded for a loaded sensor incubated in PBS. Data analysis was carried out using Octet software, version 9.0. Experimental data were fitted globally with a 1:1 Langmuir model of binding for all the antigens.

### Immunogenicity and Protection Studies Mice

Female 6- to 8-week old B6(Cg)-Tyrc-2J/J albino mice were obtained from The Jackson Laboratory. All animals were cared for in accordance with American Association for Accreditation of Laboratory Animal Care standards in accredited facilities.

### Immunizations

Vaccine immunogens were diluted in sterile PBS to their respective dose and mixed with 33.3 µL ALFQ, which, like AS01, comprises a liposomal adjuvant formulation containing monophosphoryl lipid A and QS-21 in a final volume of 50 µL. Immunization was performed intramuscularly in the quadriceps at weeks 0 and 4. Serum sampling was performed 2 weeks following the 1st and 2nd immunizations. Challenges were performed 4 weeks following the last immunization.

### IV challenge and quantification of protection

Transgenic P. berghei (strain ANKA 676m1c11, MRA-868) expressing full-length P. falciparum CSP and a green fluorescent protein/luciferase fusion protein (Pb-PfCSP-GFP/Luc-SPZ) were obtained. Approximately 4 weeks following the final immunization, mice were intravenously challenged in the tail vein with 2000 freshly harvested Pb-PfCSP-GFP/Luc-SPZ in Leibovitz’s L-15 medium (Thermo Fisher Scientific Inc., Waltham, MA, USA). Then, 40–42 h post-challenge, mice were injected intraperitoneally with 150 μL of D-luciferin (PerkinElmer, Waltham, MA, USA; 30 mg/mL), anesthetized with isoflurane, and imaged with the IVIS® Spectrum in vivo imaging system (PerkinElmer, Waltham, MA, USA) 10 min after luciferin injection. Liver burden was quantified by analyzing a region of interest (ROI) in the upper abdominal region; the total flux (p/s) was measured using the manufacturer’s software (Living Image 4.5, PerkinElmer, Waltham, MA, USA). Percent (%) protection for each mouse was calculated based on the liver burden flux (p/s) data. Percent (%) protection = [100 − ((antibody-treated mouse flux/geometric mean flux of untreated mice) × 100)].

### Serology

Immune responses were measured to full-length PfCSP or to 15-mer peptides by Biolayer interferometry (BLI). Biotinylated flCSP and peptides were captured on streptavidin (SA) biosensor and dipped into 100-fold diluted serum samples from individual mice. Assays were performed with agitation at 30°C and the binding response was calculated. In all Octet measurements, parallel correction to subtract systematic baseline drift was carried out by subtracting the measurements recorded for a loaded sensor incubated in PBS. Data analysis was carried out using Octet software, version 9.0.

### Adoptive Transfer and Immunization

B cells were purified from iGL_CIS43 BCR knock-in CD45.2 mice using the Pan B Cell Isolation Kit II (Miltenyi Biotec). The cells underwent staining with BCR specific probes and surface-staining antibodies, followed by FACS analysis (BD FACSymphony S6) to determine precursor frequency. Afterward, the cells were suspended in PBS to a total of 200 µl and administered to B6.SJL-Ptprca Pepcb/BoyJ host mice (CD45.1) via tail vein injection. Immunogens were prepared by diluting them in PBS to a final volume of 100 µl, then combined with 2% Alhydrogel adjuvant (Invivogen) in a 1:1 ratio to reach a total volume of 200 µl. This mixture was incubated on a rotator for 30 minutes and administered intraperitoneally to the host mice 24 hours post-adoptive transfer. All procedures involving animals were performed following protocols approved by the Institutional Animal Care and Use Committee (IACUC) of Harvard University and Massachusetts General Brigham.

### Flow Cytometry

Spleens from immunized mice were collected and mechanically disrupted using a 70 µm filter to isolate single-cell suspensions. Red blood cells were lysed with ACK buffer, and the remaining cells were washed in PBS supplemented with 2% FBS (FACS buffer). The cell suspensions were incubated with Live/Dead Blue dye and an Fc receptor blocking reagent in FACS buffer at 4°C for 10 minutes, followed by washing. Biotinylated probes conjugated with fluorochrome-streptavidin at a 4:1 molar ratio were prepared at 4°C for 20 minutes and diluted to a final concentration of 50 nM in FACS buffer. These were incubated with the cells at 4°C for 30 minutes. Surface-staining antibodies were diluted in FACS buffer at ratios of 1:100 to 1:400 and added to the cells for 30 minutes at 4°C. After two washes, samples were either analyzed or sorted using the BD FACSymphony S6 system. Single cells were dry-sorted into 96-well PCR plates, snap-frozen on dry ice, and stored at −80°C for subsequent sequencing. FACS data were analyzed using FlowJo version 10.

### B cell sorting

Mouse splenocytes were collected, resuspended in 90% Fetal Bovine Serum (FBS)/ 10% DMSO at ∼107 cells/ml, and stored in liquid nitrogen as previously described. On day of sorting, cryopreserved splenocytes were thawed in 37°C water bath, spun down at 500 g for 5 min, and resuspended in 1 ml of PBS. Spun down the cells again, removed PBS, and resuspended them in 1:1000 diluted VIVID in PBS and incubated on ice for 15min in dark. In the meantime prepared a staining mix in PBS/2% FBS containing BUV395-labeled anti-mouse CD3, CD4 and CD8 antibodies (T cell markers), PerCP.Cy5.5-labelled anti-mouse F4/80 (monocyte marker), Texas Red-labeled anti-B220 (B cell marker), BV711-labeled anti-mouse IgD and anti-mouse IgM, FITC-labelled anti-mouse IgG1, IgG2a, IgG2b and IgG3 antibodies, PE-labeled PfCSP, PECy7-labeled P29 and APC-labeled P21. After washing off the VIVID stain with 1 ml of cold PBS, resuspended each splenocyte sample in 200 μl of staining mix and stained on ice and in dark for 30 min. Washed the cells 2x with cold PBS/2% FBS (1ml each time) and filtered out cell aggregates with a 40 μm cell strainer. The stained splenocytes were sorted on a BD Symphony 6 sorter. PfCSP+/P29+/P21+ IgG+/IgD-/IgM-B cells were single-sorted to a 96-well plate containing 5 μl /well of cell lysis buffer (Qiagen TCL with 30 μl of β-mercaptoethanol per ml), snap-frozen on dry ice and stored at −80°C for RATPIg processing later.

### Rapid Amplification-Transfection-Production of Immunoglobulins (RATPIg)

The sorted B cells were subjected to RATP-Ig following the procedures as described previously^51^. Briefly, single-cell RNA was purified with RNAclean beads (Beckman Coulter). cDNA was then synthesized using 5′ RACE reverse-transcription and amplified by PCR, and heavy and light chain variable regions enriched. An aliquot of enriched cDNA was sequenced using 2 × 150 paired-end reads on an Illumina MiSeq. For immunoglubulin production, enriched variable regions were assembled into expression cassettes that include CMV, and HC/LC-TBGH polyA fragments. Assembled cassettes were amplified by PCR and transfected into Expi293 cells in 96-well deep-well plates using the Expi293 Transfection Kit (ThermoFisher Scientific). Cell cultures were grown at 37°C, 8% CO2, and 1100 RPM shaking for 5–7 days. Cell culture supernatants were harvested by centrifugation.

### ELISA screening of RATPIg supernatants

Streptavidin-coated 96 well plates (Pierce) were coated with 100 μl/well of 2 μg/ml biotinylated P21 or P29 and high binding costar 96 well plates were coated with 100 μl/well of 2 μg/ml PfCSP at 4°C overnight. The coated plated were washed once with PBS/T (PBS/ 0.05% Tween-20) and blocked with 1:10 diluted ImmuneTech Blocking solution for 1 hr at room temperature. After 3x PBS/T wash of the ELISA plates, 100 μl of RATPIg supernatants in 96-well deep well plates were transferred to the antigen coated ELISA plates and incubated at room temperature for 1 hr. The ELISA plates were washed 5x with PBS/T and incubated with 100 μl of 1:5,000 diluted peroxidase-conjugated goat-anti-mouse IgG at room temperature for 1 hr. After 5x PBS/T wash, 100 μl of SureBlue™ TMB 1-Component Microwell Peroxidase Substrate was added to each well of the ELISA plate and incubated for 10 min at room temperature. The colorogenic reaction was terminated with 100 μl /well of 1N Sulfuric Acid and the plates were read for OD450 on a SpectraMax Plus microplate reader (Molecular Device).

### Statistical analysis

Statistical analyses were performed using two-tailed Mann-Whitney tests with GraphPad Prism 10.2.3 software (La Jolla, CA). Differences were considered statistically significant at *P* ˂ 0.05.

## DATA AVAILABILITY

All other data from the current study are available from the corresponding authors on reasonable request.

## ACKNOWLEDGEMENTS

We thank A.V.S. Hill for the R21 vaccine, J. Stuckey for assistance with figures, Walter Reed Army Institute of Research (WRAIR) for ALF-Q adjuvant, members of the Flow Cytometry Core at the Ragon Institute for assistance, and members of the Vaccine Research Center, NIAID, NIH, for discussions and comments on the manuscript. This research was supported, in part, by the Intramural Research Program of the National Institutes of Health (NIH). The contributions of the NIH authors are considered Works of the United States Government. The findings and conclusions presented in this paper are those of the authors and do not necessarily reflect the views of the NIH or the U.S. Department of Health and Human Services. This research was, in part, supported by the National Cancer Institute’s National Cryo-EM Facility at the Frederick National Laboratory for Cancer Research under contract HSSN261200800001E. This work was also supported by the NIH NIAID R01 AI168114 and R01 AI151178; Leidos Biomedical Research Inc. No. 20X013F; the National Research Foundation of Korea’s Basic Science Research Program NRF-2021R1A6A3A14044219; and the Ragon Institute of Mass General Brigham, MIT, and Harvard.

## AUTHOR CONTRIBUTIONS

The overall design of the study was by P.T., B.Z., A.H.I., R.A.S. and P.D.K; Experiments and analyses were performed by P.T., L.D.S.P., M.D., X.C., J.H.K, K.N., B.Z., M.L., A.R., S.K., W.P.K., T.S.; Nanoparticle immunogens were produced and characterized by P.T.; Animal studies were performed by L.D.S.P, M.D., M.L.; Antigenicity and serology was performed by P.T.; KI-mice immunization and associated analyses were done by J.H.K; B-cell sorting and ELISA was performed by X.C., K.N., and W.P.K.;; RATPIg was done by A.R., and S.K.; Nanoparticle designs were made by B.Z., P.T., and P.D.K. Negative stain electron microscopy by done by P.T., T.S., and Y.T.; D.C.D. supervised RATPIg sequencing; F.D.B. supervised KI-mice immunization; A.H.I. and R.A.S. supervised malaria challenge studies; P.D.K., T.Z. oversaw the project and wrote the manuscript along with P.T., J.H.K., X.J., with input and comments from all authors.

