## Supplemental Information for "Protective immunity against malaria by a nanoparticle CIS43-based junctional vaccine alone or in combination with R21"

**Table S1**

| **Construct name**<br><br>**(Nterm attachment-nanoparticle-number of repeats-Cterm attachment)** | **Amino acid sequence** |
|---|---|
| **NPDP19-LuS-PADRE**<br>N-term attachment – NPDP19<br>Nanoparticle- LuS<br>Number of repeats – 1<br>C-term attachment – PADRE | mdskgssqkgsrllllvvsnlllpqgvvgKQPADGNPDPNANPNVDPNgsgsaMQIYEGKLTAEGLRFGIVASRFNHALVDRLVEGAIDAIVRHGGREEDITLVRVPGSWEIPVAAGELARKENISAVIAIGVLIRGATPHFDYIASEVSKGLADLSLELRKPITFGVITADTLEQAIERAGTKHGNKGWEAALSAIEMANLFKSLRGAKFVAAWTLKAAA |
| **2T-LuS-PADRE**<br>N-term attachment – NPDP19<br>Nanoparticle- LuS<br>Number of repeats – 2 in tandem<br>C-term attachment – PADRE | mdskgssqkgsrllllvvsnlllpqgvvgKQPADGNPDPNANPNVDPNggsggKQPADGNPDPNANPNVDPNgsgsaMQIYEGKLTAEGLRFGIVASRFNHALVDRLVEGAIDAIVRHGGREEDITLVRVPGSWEIPVAAGELARKENISAVIAIGVLIRGATPHFDYIASEVSKGLADLSLELRKPITFGVITADTLEQAIERAGTKHGNKGWEAALSAIEMANLFKSLRGAKFVAAWTLKAAA |
| **PADRE-LuS-2T**<br>N-term attachment – PADRE<br>Nanoparticle- LuS<br>Number of repeats – 2 in tandem<br>C-term attachment – NPDP19 | mdskgssqkgsrllllvvsnlllpqgvvgAKFVAAWTLKAAASLVRgsgsaMQIYEGKLTAEGLRFGIVASRFNHALVDRLVEGAIDAIVRHGGREEDITLVRVPGSWEIPVAAGELARKENISAVIAIGVLIRGATPHFDYIASEVSKGLADLSLELRKPITFGVITADTLEQAIERAGTKHGNKGWEAALSAIEMANLFKSLRGggsggKQPADGNPDPNANPNVDPNggsggKQPADGNPDPNANPNVDPN |
| **3T-LuS-PADRE**<br>N-term attachment – NPDP19<br>Nanoparticle- LuS<br>Number of repeats – 3 in tandem<br>C-term attachment – PADRE | mdskgssqkgsrllllvvsnlllpqgvvgKQPADGNPDPNANPNVDPNggsggKQPADGNPDPNANPNVDPNggsggKQPADGNPDPNANPNVDPNgsgsaMQIYEGKLTAEGLRFGIVASRFNHALVDRLVEGAIDAIVRHGGREEDITLVRVPGSWEIPVAAGELARKENISAVIAIGVLIRGATPHFDYIASEVSKGLADLSLELRKPITFGVITADTLEQAIERAGTKHGNKGWEAALSAIEMANLFKSLRGAKFVAAWTLKAAA |
| **PADRE-LuS-3T**<br>N-term attachment – PADRE<br>Nanoparticle- LuS<br>Number of repeats – 3 in tandem<br>C-term attachment – NPDP19 | mdskgssqkgsrllllvvsnlllpqgvvgAKFVAAWTLKAAASLVRgsgsaMQIYEGKLTAEGLRFGIVASRFNHALVDRLVEGAIDAIVRHGGREEDITLVRVPGSWEIPVAAGELARKENISAVIAIGVLIRGATPHFDYIASEVSKGLADLSLELRKPITFGVITADTLEQAIERAGTKHGNKGWEAALSAIEMANLFKSLRGggsggKQPADGNPDPNANPNVDPNggsggKQPADGNPDPNANPNVDPNggsggKQPADGNPDPNANPNVDPN |

* Red: signal sequence; blue: PADRE; orange: cleavage site; purple: linker

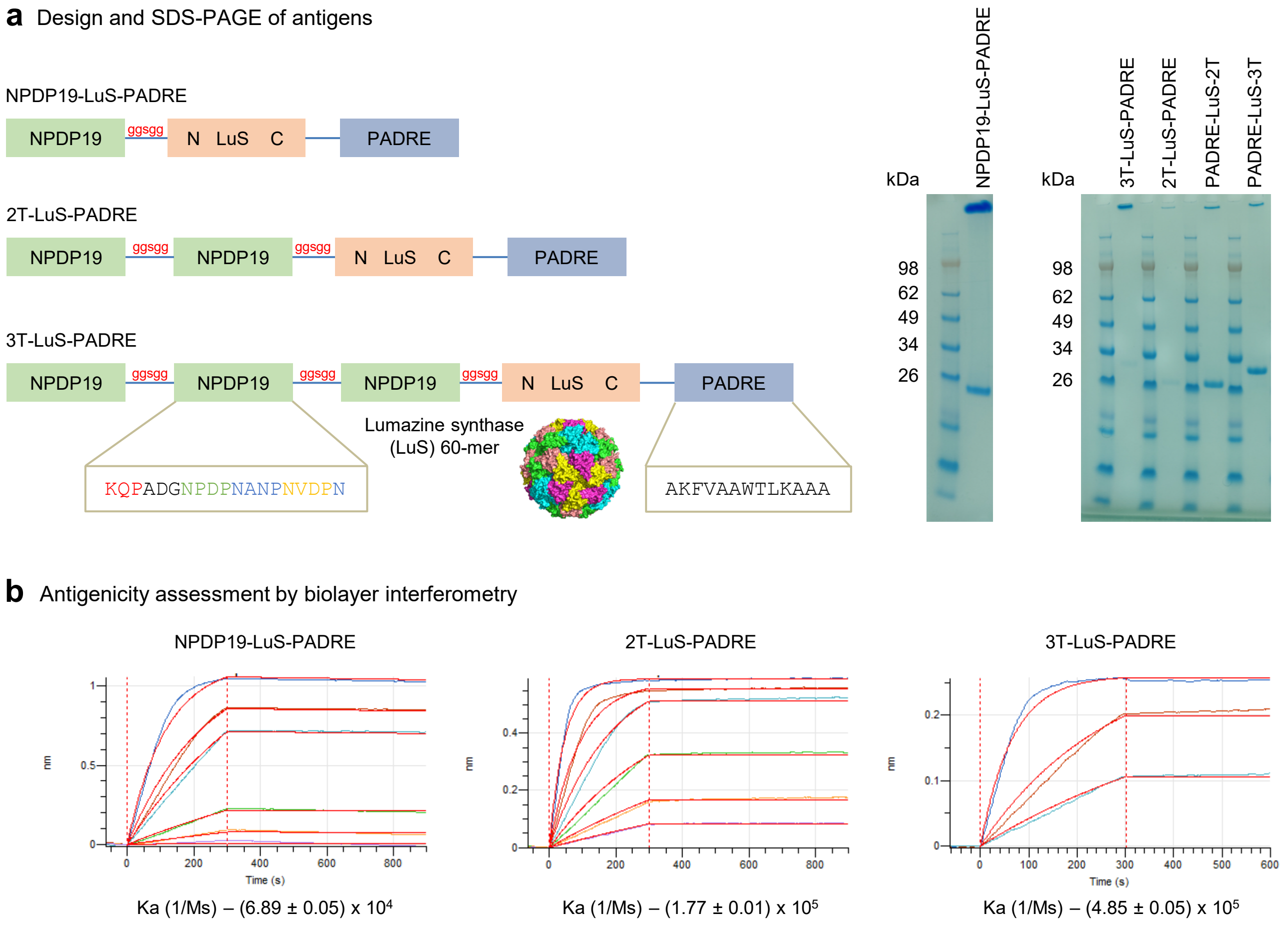


**Supplementary Figure 1. Characterization of immunogens by SDS-PAGE and biolayer interferometry related to Fig 1.**

**a** Design schematic and SDS-PAGE of antigens. **b** Binding assessment of antigens to antibody D3 by biolayer interferometry.

Correlation analyses of antibody titers and protection

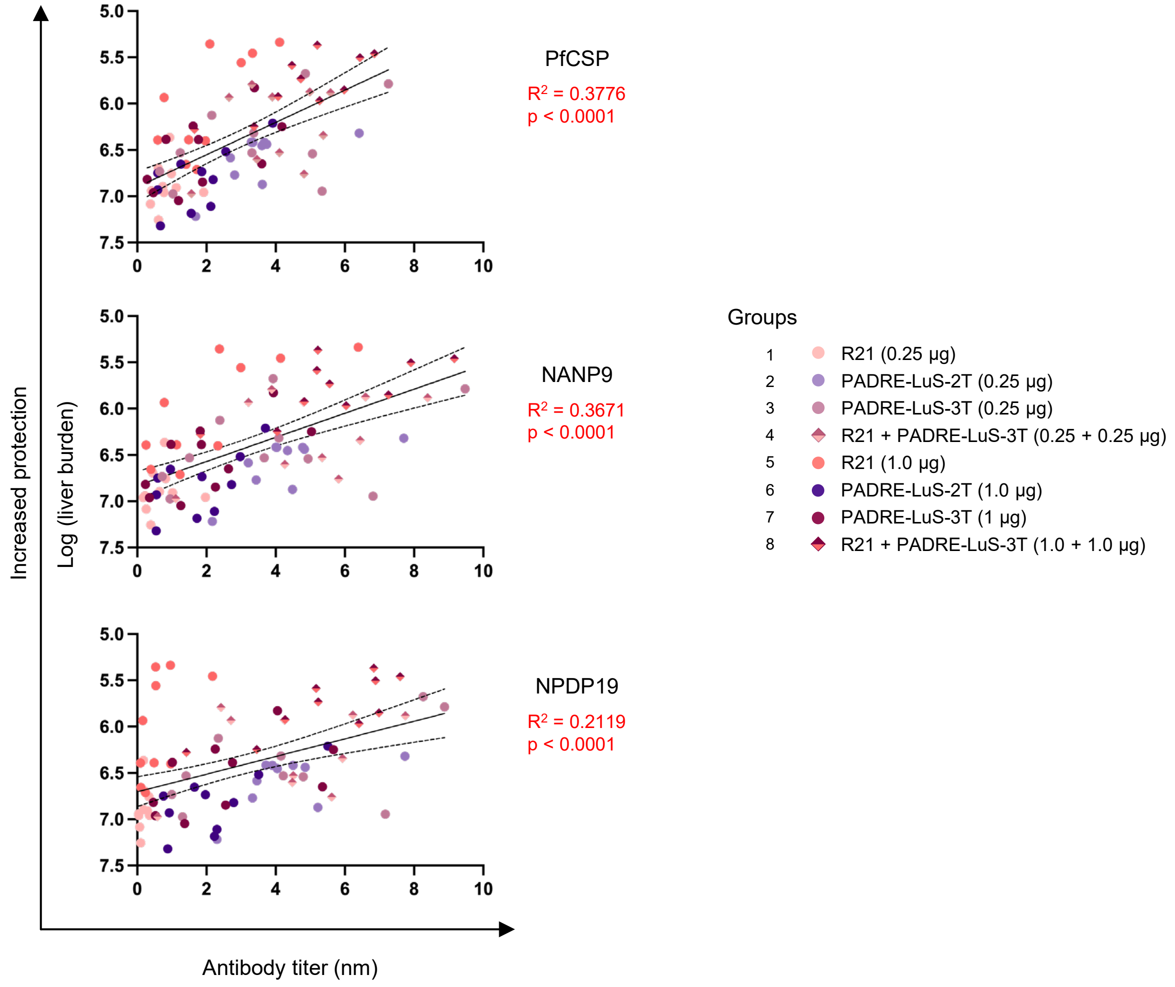


**Supplementary Figure 2. Protective efficacy and antibody titers correlation, related to Fig 4.**
Correlation analyses for week 6 antibody titers and liver burden (total flux) for three probes; PfCSP, NANP9 (major), and NPDP19 (junction). Simple linear regression analysis was performed in GraphPad Prism version 9.0.

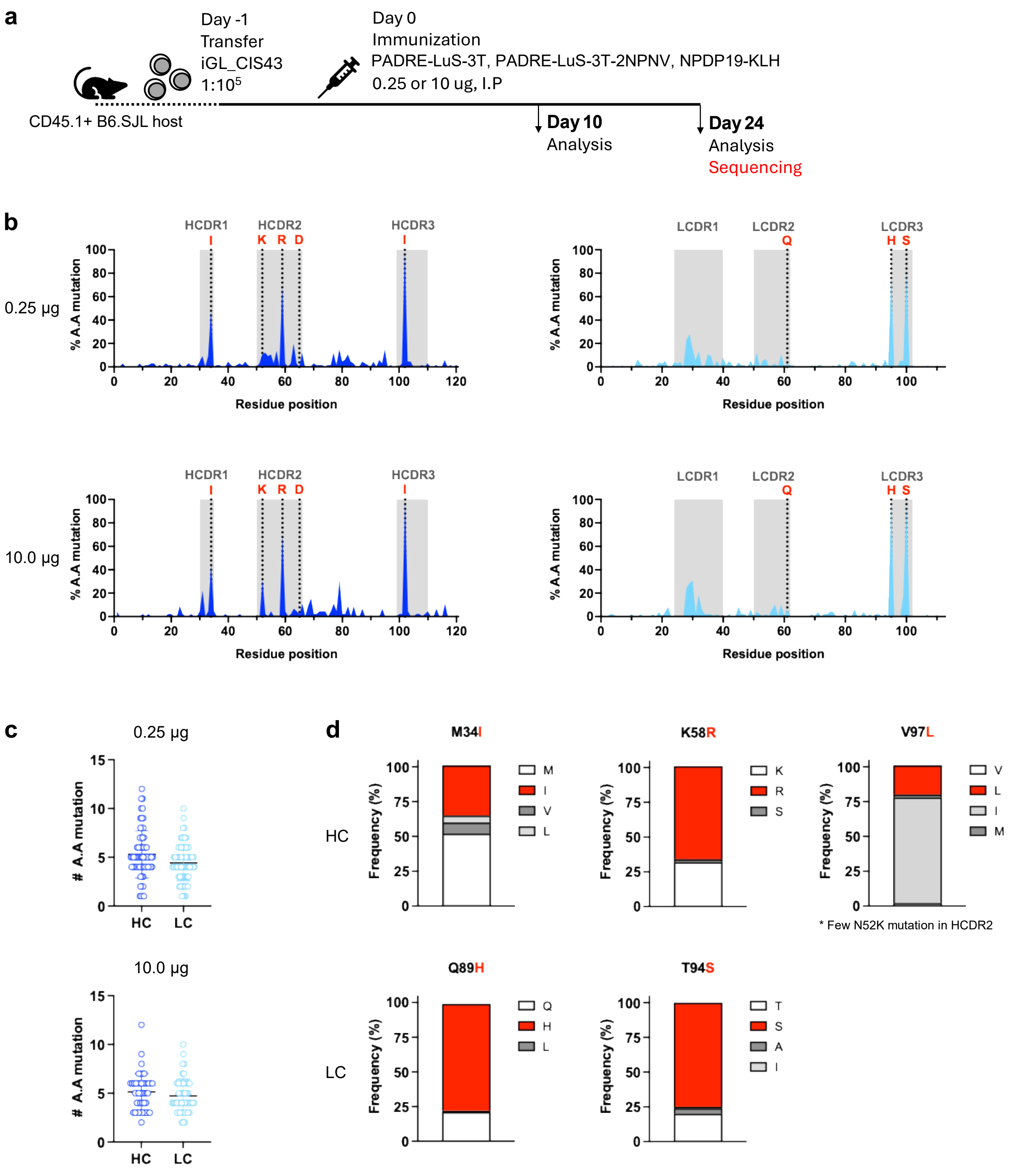


**Supplementary Figure 3. Mutational frequency from iGL-CIS43 mice immunized with PADRE-LuS-3T, related to Fig. 5.**
**a** Schematic of adoptive transfer model. **b** Hotspot analysis shows frequency of observed HC and LC mutations per residue at 24 DPI. HCDRs and LCDRs are highlighted in gray. Letters in red (only present in mature CIS43 HC) indicate key aa residues for the recognition of the junctional epitope. **c** Total amino acid mutations for HC and LC are shown. **d** Distribution frequency of select iGL-CIS43 B cell HC and LC aa mutations at positions 34, 58, 97, 89, and 94.

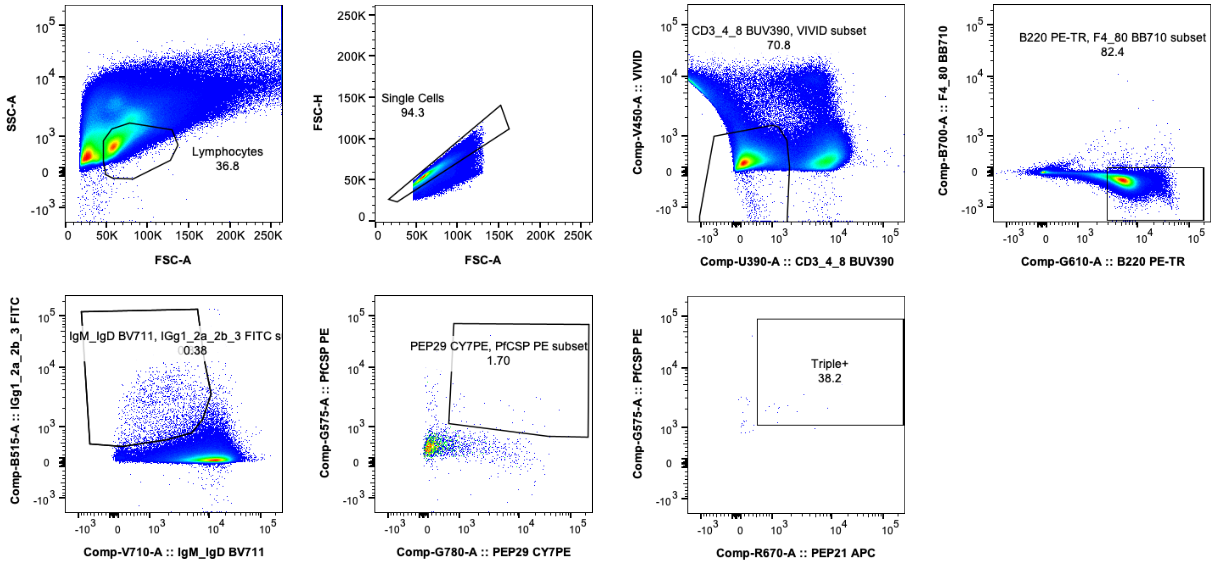


**Supplementary Figure 4. B-cell sorting data, related to Fig. 6.**
B-cell sorting steps to collect PfCSP/P21/P29 triple positive IgG+ B cells for RATPIg. P21 (junction); P29 (major repeat).
